# Where Initial rates are directly proportional to Substrate concentrations with Application in Molar-mass Determination, Zero-order Specificity constant is Inappropriate

**DOI:** 10.1101/2023.04.06.535898

**Authors:** Ikechukwu Iloh Udema

## Abstract

**Background:** “High-ranking scientists” employ the initial rate (*v*_i_), expression without consideration for the conditions under which the *v*_i_ expression can be used. The consequence is the suggestion that the *v*_i_ is equal to the product of maximum velocity, *V_max_, and* substrate concentration [*S*_0_] divided by the Michaelis-Menten constant, *K*_M_.

**Objectives:** The main objectives are: 1) to show that *v*_i_ is not equal to *V*_max_[*S*_0_]/*K*_M_; 2) to show that the equilibrium dissociation constant, *K*_d_, is strictly proportional to the concentration ([*E*_0_]) of the enzyme; and 3) to show that the two standard quasi-steady-state assumptions (sQSSA) and reverse QSSA (rQSSA) have a limited domain of validity.

**Methods:** The study was experimental and theoretical, supported by the Bernfeld method of enzyme assay.

**Result:** *K*_d_ is directly proportional to [*E*_0_], and *v*_i_ is not equal to *V*_max_[*S*_0_]/*K*_M._. A *K*_M_-like value that is greater than the putative *K*_d_ value, 2.482 g/L, is equal to 2.569 g/L. The *K*_M_-like values in other situations are 2.396 and 2.407 g/L; the corresponding equilibrium dissociation constant (*K*_d_) values are, respectively, 2.288 and 2.299 g/L; the molar mass of insoluble potato starch ranges between 62.296 and 65.616 exp. (+6) g/mol.

**Conclusion:** The equations that invalidate the assumption that *v*_i_ is equal to *V*_max_[*S*_0_]/*K*_M_ whenever [*S*_0_] is much less than *K*_M_ were derived; the proposition that *K*_d_ is strictly proportional to [*E*_0_] was confirmed; the molar mass of starch could be calculated from the derived equation; and it was shown graphically and mathematically that both the sQSSA and rQSSA domains have a limit of validity; the equation with which to calculate the second order rate constant based on the conditions that validate the rQSSA is not applicable to the sQSSA. A *K*_M_-like value that is greater than the putative *K*_d_ value is possible.

**GRAPHICAL ABSTRACT:** The graphical abstract illustrates three zones: the zone in which the sQSSA is valid, the zone in which the rQSSA is valid, and the zone in which neither assumption is exclusively valid. The curved arrow (oxblood) pointing to the red line depicts a tendency towards conditions that validate the rQSSA if the assay is conducted with an appropriate [S_0_]/[E_0_] ratio (< 1 to ≪1) while the red curved arrow pointing to the blue line depicts a tendency towards conditions that validate the sQSSA if the assay is conducted with an appropriate [S_0_]/[E_0_] ratio (>1 to ≫1). The enzyme-substrate complex (ES) is in a quasi-steady state with respect to S as depicted by ∂ [ES]/ ∂t≈0, the sQSSA case, while in the rQSSA, it is the S that is in a quasi-steady state with respect to ES as depicted by ∂[S_0_]/∂t≈0. The double-headed arrow merely shows, artistically, the limit of the data points.

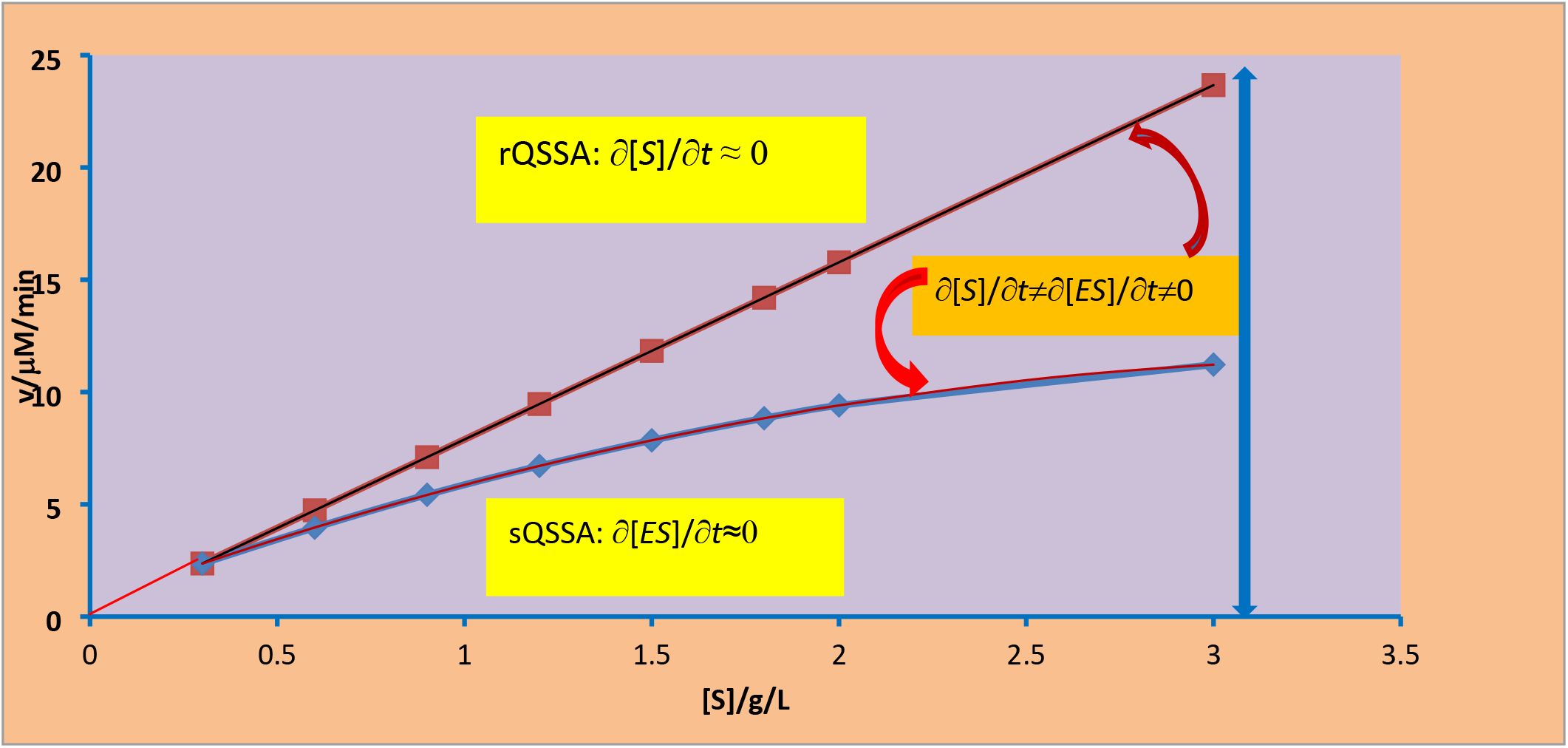

## 1. INTRODUCTION

There are remarkable studies on the problem associated with the generation of valid kinetic parameters anchored on the conditions that validate certain assumptions; such assumptions are the well-known constancy of the rate of product, P, formation, such that d[*ES*]/d*t* (or Δ[*ES*]/Δ*t*) is ≈ 0 and d[*S*_0_]/d*t*(or Δ[*S*_0_]/Δ*t*) is ≈ 0, where [ES] and [*S*_0_] are the concentration of the enzyme-substrate complex and the initial concentration of the substrate. A question arising from studies is: Are there parameter domains where instead of ES (i.e., the molar concentration of the enzyme, E, which formed a complex with the substrate, S) being in a quasi-steady state with respect to S, there is a “reverse quasi-steady-state (rQSSA)” in which S is in a quasi-steady state with respect to ES? [1] The question implies that there is unarguably the opposite assumption: the standard QSSA (sQSSA) that seems to have been originally credited to Briggs and Haldane [2] and Savageau [3]. The rQSSA is also regarded as an alternative definition of quasi-steady state [4], where, as above, d[*S*_0_]/d*t* ≈ 0; this was originally attributed to Segel and Slemrod [1]. Within this approximation at high enzyme concentration [*E*_0_], the conditions [4] whereby [*E*_o_] is ≫ [*S*_0_] and [*E*_o_] is ≫*K* (where *K*, the Van Slyke–Culen constant [5], is =*k*_cat_/*k*_1_, where *k*_cat_ and *k*_1_ are the catalytic rate of product formation and the second order rate constant for the formation of ES, respectively), were used for the derivation of appropriate equations as originally attributed to Schnell and Maini [6]. This notwithstanding, there is a view that the condition whereby [*S*_0_] is ≫ [*E*_o_] is unnecessarily restrictive [7]. As such, the Michaelis–Menten equation can be used even when [*S*_0_] is ≈ [*E*_0_] as long as the Michaelis–Menten constant, *K*_M_, is ≫ 1 or [*E*_0_] is ≪ *K*_M_ [7]. Consequently, sQSS can also be valid without such restrictions. The argument in this research is that despite the conditions that minimise the restriction on the parameter domains for which sQSSA and the Michaelis-Menten equation remain valid, there is, after all, a limit to such domains.

Against the backdrop of the facts and principles enunciated above, it has become necessary to support the view that the velocity (initial rates, *v*_i_) equations of the catalytic reaction have been employed for the determination of kinetic parameters on a number of occasions outside of the conditions for which they are valid [8]. This implies that some kinetic parameters, including the specificity constants, may not have qualified as sQSSA, rQSSA, *etc*. As in an *in vivo* scenario, [*E*_0_] may be ≫ [*S*_0_], or the former may be approximately of the same order of magnitude as its substrate concentration [6]. The former scenario ([*E*_0_] ≫ [*S*_0_]) should be in line with rQSSA, while the latter ([*E*_0_] ≈ [*S*_0_]) scenario may be partially in line with sQSSA. The goal of the study is to show that it is the rQSSA that is applicable where initial rates are directly proportional to the concentration of the substrate, apart from the usual observation that [*E*_o_] must be ≫ [*S*_0_]. Under such a condition, it is possible to show that the molar mass of the substrate, such as starch, can be determined. By so doing, one can reveal that where initial rates are directly proportional to substrate concentrations with application, in substrate molar-mass determination, zero-order specificity constant is inappropriate. This can be accomplished with the following objectives: 1) to derive an equation that invalidates the assumption that whenever [*S*_0_] is ≪ *K_M_* (and in particular, when [*E*_0_] is ≫ [*S*_0_]), *v*_i_ is always = *V*_max_ [*S*_0_]/*K*_M_; 2) to derive an equation that shows that the ES equilibrium dissociation constant is strictly proportional to [*E*_0_); 3) to calculate, based on the derived equation, the *K*_d_ value compared with a graphical value; 4) to apply the rQSSA-based derived equation in the calculation of the molar mass of the polymer substrate; and 5) to illustrate graphically and mathematically, a limit to the extent of the parameter domain in which the QSS and Michaelian equation can be valid.

### 1.1 Significance

For the first time, compared to the best of the available pieces of information in the literature, the hidden un-Michaelian kinetics that reduces the accuracy of Michaelian kinetic parameters has been unraveled. The error stems from the fact that the first two to three (or more) initial rates may be directly proportional to the concentration of the substrate and, separately, can yield a negative intercept in a double reciprocal plot. Such negative intercepts contribute to the less accurate values of the kinetic parameters generated by whatever means—direct (or reciprocal variant) linear plot, double reciprocal plot, nonlinear regression analysis, *etc*. The derivations have enabled the determination of the ES dissociation constant and the molar mass of the substrate, starch, in this study based on a kinetic model.

## 2.0 THEORY

If the initial rates are directly proportional to the concentrations of the substrate, the coefficient of determination is very likely to be ≥ 0.999 (it could be =1); being < 0.999 may be as a result of error in measurement of initial rates, initial substrate concentration, timing, *etc.*, leading to “outliers” as often referred to in the old literature [9, 10] and in papers [9, 11–13] devoted to how best to produce accurate initial rates (*v*_i_) or rather kinetic parameters following the assay of an enzyme. The notion that *v*_i_ is directly proportional to [*S*_0_], where the proportionality constant is the ratio of the maximum velocity of enzymatic action to the Michaelis-Menten constant (or the zero-order specificity constant), can be found in many standard undergraduate text books and in high-ranking journals containing views about the *in vivo* concentration of the enzyme compared with the substrate [6, 8].

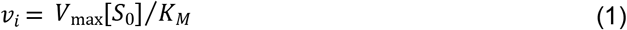

where *V*_max_ and *K*_M_ are the maximum velocity of catalytic action and Michaelis-Menten constant respectively. Equation (1) stems from the fact that, in certain situations, the concentrations of the substrate are ≪ *K*_M_ and the concentration of the enzyme [*E*_0_] could be ≫ [*S*_0_] as is the case in an *in vivo* scenario [6, 8]. “One question that needs an answer is: does it mean that after the consumption of 4-6 slices of bread, the concentration of the enzyme in the small intestine is > the overall concentration of a carbohydrate-rich diet”? Equation (1) originates from the Michaelis-Menten equation given below:

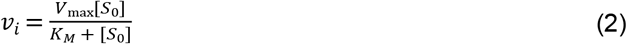

Thus, if, by conceptual and operational arguments, the enzyme-catalysed reaction cannot attain half its maximum rate of catalysis until substrate concentration equal to the *K*_M_ is available, it should be inappropriate to convert Eq. (2) to Eq. (1).

It falls within the realm of common sense to observe that if [*E*_0_]*(1)* is > [*E*_0_](2), *K*_M_ for the former should be proportionately < the *K*_M_ for the latter. It has been observed in the literature that a high-ranking biochemist [14], whose authority in the field is almost the kind no one dares question, has consistently called for the direct measurement of specificity constant (*V*_max_/*K*_M_); thus, dividing the latter obtained from the plot of *v*_i_ versus [*S*_0_] by [*E*_0_] should translate into the direct measurement of specificity constant (SC), even if [*S*_0_] is < [*E*_0_]. This is definitely inappropriate. Arguments about the appropriateness of SC as defined in Eq. (1) will be subject to some aspects of the quasi-steady-state assumption in due course.

If, indeed, *v*_i_ is directly proportional to [*S*_0_], then the following relationship should hold.

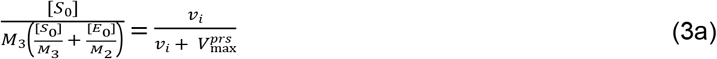

where, *M*_3_, *M*_2_ are the pre-steady-state maximum velocity (PRSV), molar mass of substrate, and molar mass of the enzyme, respectively. Issues regarding PRSV have been investigated elsewhere [15] However, the choice of PRSV is purely a coincidence; otherwise, the equations being derived are equally applicable to any linear phase, including early steady-state (SS) [16].

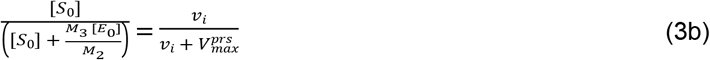

Expanding Eq. (3b) gives:

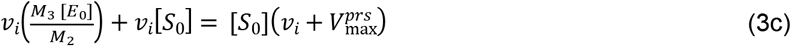

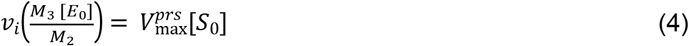

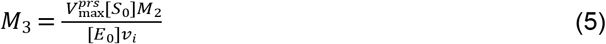

An unbiased critical examination of Eq. (5) shows that, the equation can only be valid if *v*_i_ is totally and directly proportional to [*S*_0_] and, expectedly, should be a constant for a given concentration of the enzyme, which in turn influences the magnitude of *v*_i_. The coefficient of determination (*R*^2^) could be = 1. Thus, Eq. (5) falls outside the realm of Michaelian kinetics or the quasi-steady-state approximation (QSSA). It should be applicable to reverse QSSA (rQSSA). This is the core reason why the PRSV or its SSV counterpart is a better choice because it is much lower than the zero-order (asymptotic) values of the actual maximum velocity.

The second equation is:

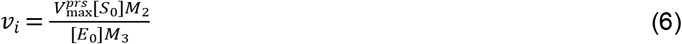

Equation (6) clearly shows that the proportionality constant, φ (or slope) is given as:

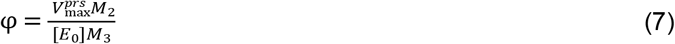

Either from Eq. (6) or Eq. (7), the most important observation is that, the reciprocal of the equilibrium dissociation constant (*K*_d_), otherwise called the association constant (*K*_a_), is given as:

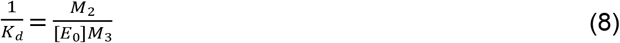

Based on Eq. (8), one can convincingly opine that like *K*_M_, *K*_d_ is directly proportional to [*E*_0_]. However, the enabling scenarios differ for *K*_M_ and *K*_d_; while the [*S*_0_] range for the former must fall between values < the *K*_M_ and values ≫ *K*_M_, the [*S*_o_] range for the latter must be ≪ *K*_M_ if known *a priori.* Most importantly, the [*E*_o_] value suitable for *K*_M_ must be ≪ [*S*_0_] in line with expectations of standard QSSA (sQSSA), while, in addition, [*E*_0_] must be ≫ [*S*_0_] for the determination of *K*_d_ as expected in a reverse QSSA (rQSSA) scenario. It follows from Eq. (8) that if the values of *M*_2_ and *M*_3_ are known, the *K*_d_ for any enzyme of known concentration can be calculated and used to estimate the [*S*_0_] range that falls between values < *K*_d_ and values ≪ *K*_M_.

With this background theory, it is clear that there are cogent reasons to rewrite Eq. (1), which becomes:

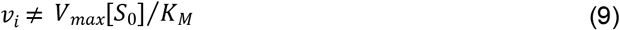

Thus, in place of Eqs (1) and (9), the following equation applies:

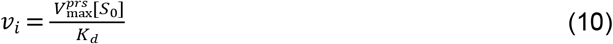

There is hardly any one-substrate-one-enzyme reaction in which the reverse reaction and forward reaction may not occur. The difference lies in the magnitude of *V_max_,* which may be high if [*E*_T_] is high with either the correspondingly much higher values of [*S*_T_] for the sQSSA case or much lower [*S*_T_] values for the rQSSA case; thus the following may hold:

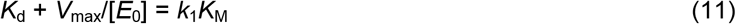

Equation (11) is therefore, strictly speaking, not applicable to rQSSA, but rather it is applicable to sQSSA.

Equation (11) is despite the view that where [*E*_0_] ≫ [*S*_T_], the following equation, which is very similar to the Michaelis-Menten equation, is applicable.

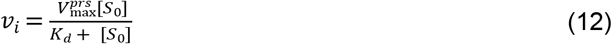

In this research, however, Eq. (12) is redesignated as one that is appropriate for a situation in which [*E*_0_] is ≈ [*S*_0_]. This is to imply that any plot of *v*_i_ versus [*S*_0_] may not be far from Michaelian kinetics, but the zero-order (or asymptotic) value of the maximum velocity is not attainable under such a situation. This is in line with the view elsewhere [17] that “when both the sQSSA and rQSSA are invalid, the initial enzyme and substrate concentrations are comparable”. A double reciprocal linearisation of Eq. (12) gives the slope as: 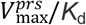, yet, 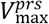 is < the magnitude appropriate for [*E*_0_] if assayed with a substrate concentration range that does not include saturating concentrations of the substrate. Because Eq. (11) is more relevant to the Michaelian equation, Eq. (12) can be intuitively related to Eq. (10) as follows:

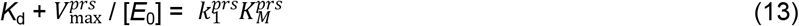

Equation (13) is born out of a reasonable postulation to the effect that:

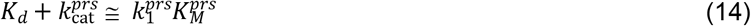

where and are respectively the 2^nd^ order rate constant, which is > the pre-steady-state value but < the zero-order value, and the Michaelis-Menten-like constant, which is > *K*_d_ but < *K*_M_. This simply means that *K*_d_ in Eq. (12) may be replaced by a parameter that is therefore neither the true *K*_d_ nor the true *K*_M_. Again, this implies that it is only a situation where the enzyme attains total saturation that guarantees the true value of a *K*_d_, which may be equal to the value as defined by Eq. (8). Also, given different values of substrate concentration range, for the same concentration of the enzyme, different values of *K*_d_ are expected.

### 2.1 Validity of various QSSA *vis-à-vis* appropriate definitions and values of *K_d_* and *K*_M_

The goal of this section is to examine what validates various QSSA and relate such to the values of *K*_M_ and *K*_d_ considering their definitions; this could enhance the validity of the kinetic constants that may be determined. Facts and principles justifying the preceding analysis and derivation are elucidated based on a research paper by [17] as follows: Beginning from the idea of a general equation of initial velocity [17], one writes

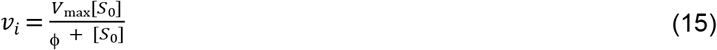

where, ϕ is given as:

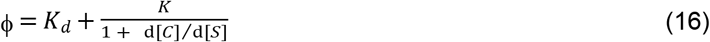

First, one considers the condition that the sum of the initial substrate concentration ([*S*_0_]) and ***K_M_*** greatly exceeds the initial enzyme concentration ([*E*_0_]), that is (but [*S*_0_] alone could be ≫ [*E*_0_]),

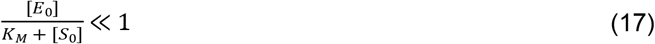

Setting d[*ES*]/d*t* ≈ 0 implies that d[*ES*]/d[S] → 0 and f =*K*_M_ in the sQSSA velocity equation. This investigation presents an equation and the possibility that it is not in compliance with the conditions that validate sQSSA. Therefore, Eqs (8) and (10) bear no iota of conceptual relevance to the equation based on sQSSA. This is apparently the reason why Borghans *et al.* [18] admitted that inequality, In-Eq. (17), cannot hold having observed that with very high concentrations of the enzyme, “*K*_M_” is small; reference to *K*_M_ is only as usual (as was the case in a very recent paper [15], though the general concept developed remains relevant]; otherwise it is appropriately the *K*_d_ (or under an exceptional circumstance to be looked into shortly, it may be considered as a special *K*_M_ different from the usual *K*_M_).

When *K*_d_ is the case, [*E*_0_] is ≫ [*S*_0_] ([*E*_0_]/[*S*_0_] 1), and the appropriate assumption is the rQSSA, otherwise known as the equilibrium approximation given as d[*S*_0_]/d*t* ≈ 0, the latter of which presupposes that ϕ is = *K*_d_. The view by Schnell and Maini [17] that “when both the sQSSA and rQSSA are invalid, the initial enzyme and substrate concentrations are comparable” is very instructive in that it goes to show that the equation derived in this study is very appropriate, being an equation in which [*E*_0_]/[*S*_0_] > 1 and not when [*E*_0_] is ≈ [*S*_0_] or of comparable magnitude; this is with reference to Eqs (3a) through (10). On the other hand, there is no way zero-order kinetics (a non-saturating phenomenon) could be the case, even if sQSSA could have been valid if [*E*_0_] is ≈ [*S*_0_] even though current opinion seems to suggest something on the contrary [7] in support of the notion that with total substrate concentration ([Ś] = [S] + [*ES*]), the parameter domain for which it is permissible to employ the classical assumption (d[*ES*]/d*t* ≈ 0) can be extended. Much earlier view is that sQSSA can provide a good approximation even when [S_0_] ≈ [*E*_0_] as long as [*E*_0_] is small compared to *K*_M_ [1].

To achieve the goal, total QSSA (tQSSA), based on the concept of total substrate concentration, is adopted. This is in addition to an unfamiliar singular perturbation method for the aggregated variable; this enables the derivation of velocity equations of substrate hydrolysis (e.g., amylolysis where applicable) and product formation [7]. For the purpose of comprehension, the total substrate concentration ([Ś]) is an aggregated or lumped variable [7]. Again, the equations are to enable core biochemists to determine kinetic parameters under conditions in which neither the sQSSA nor the rQSSA are valid [7]. The position taken in this study is that regardless of the criteria adopted that validate any of the QSSA, the foundation upon which the Michaelian concept rests cannot be jettisoned. One needs to be circumspect in ensuring that where [*S*_0_] needs to be ≫ [*E*_0_], the appropriate assumption must be inferred just as when [*S*_0_] ≪ [*E*_0_]; in other words, it is either sQSSA (d[*ES*]/d*t* ≈ 0) or, as in this study, rQSSA (d[*S*]/d*t* ≈ 0). It may not be impossible to encounter a situation in which d[*ES*]/d[*S*] is at least ≈ 1; such a situation needs to be investigated.

Further to the problem of validity, one needs to analyse the bases of the claim in this study that the parameter domain in which QSS and the Michaelian equation are valid needs not be *ad infinitum* in favour of rQSSA. If *v*_i_ is strictly proportional to [*S*_0_] for the first 2-3 data points, a double reciprocal plot can yield a small negative intercept and a larger slope. This can be illustrated in the result and discussion sections: kinetic parameters obtained in such a situation cannot be valid, and where the 2-3 data points are part of a broader range of data points, the results—the kinetic parameters—may be less accurate. As explained in a pre-print [19], [*S*_0_]_n_ *v*_n-1_ - [*S*_0_]_n-1_*v*_n_ is = zero (*n* is the number of assays according to different substrate concentrations), and consequently the equation below is expected to give an invalid result (i.e. an infinite maximum velocity and an infinite Michaelis-Menten constant).

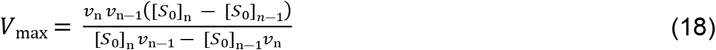

The equation for the Michaelis-Menten counterpart is:

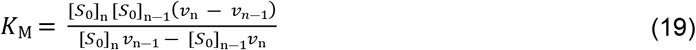

Therefore, if [*S*_0_]_n_*v*_n-1_ - [*S*_0_]_n-1_ *v*_n_ is = 0, the separate infinite values of *V*_max_ and *K*_M_ are summarily invalid, yet the specificity constant, SC, defined as *V*_max_/*K*_M_ given below, seems valid due to the absence of an infinity clause.

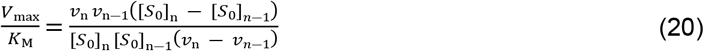

It needs to be made clear that, Eq. (20) is characteristically a general one because it is error sensitivity invariant. This is despite the fact that in the separate occurrence of the respective equations, *V*_max_ and *K*_M_ may not be valid thereby partially justifying the proposition by an imminent biochemist [14], that SC should be seen as a unique and singular kinetic parameter; it is however, very necessary to specify, the QSSA that is validly relevant to the SC generated with the assurance that substrate concentration regime (or range) matches the concentration of the enzyme in terms of either being approximately equal to, a little less than, much less than or much greater than [*E*_0_]; note that, the choice of substrate concentration range and the [*E*_0_] that validates sQSSA and Michaelian equation, does not necessarily imply that the zero-order kinetic parameters, *K*_M_, *k*_cat_ or preferably, SC were attained. So, if *v*_n_ is = 2 *v*_n-1_ and correspondingly, [*S*_0_]_n_ is 2 [*S*_0_]_n-1_, Eq. (20) transforms into:

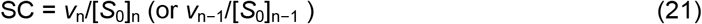

Hence, going by the definition of *V*_max_ and *K*_M_, it stands out clearly that *V*_max_/*K*_M_ is not equal to the ratio of the initial rate to the corresponding concentration of substrate, which could have been a characteristic of a single-turnover event. Hence, in circumstances in which the initial rate is consistently proportional to the concentration of the substrate (with the possibility that the coefficient of determination is ≥ 0.999), Eq. (21) represents SC for a scenario where rQSSA is valid (QED). Another equation in the literature [20] that can redefine the limit of the parameter domain for which sQSSA and rQSSA are valid is given as follows:

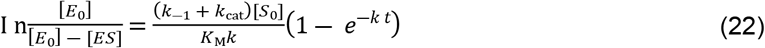

where *k*_-1_, *k*_cat_, *k*, and *t* are the reverse first-order rate constants for the dissociation of ES into free E and S, the catalytic first-order rate constant, the pseudo-first order rate constant for the utilisation of the substrate, and the duration of ES formation (or the life span) of ES.

While Eq. (22) represents a general principle in terms of what it represents, *K*_M_ in the equation may not be the actual *K*_M_ if, according to a recommendation in the literature [8], the agreement between the sQSSA solution and the numerical solution is quite good when [*E*_0_] ≤ 0.01[*S*_0_]. The simple issue is that some of the concentrations of the substrates must be about 40 to 100-fold higher than [*E*_0_]. Any substrate concentration range < 40 to 100-fold may not be in good agreement with “zero-order level” sQSSA. In such a scenario, rQSSA may be relevant and Eq. (22) may be applicable because in the equation given below (Eq. (23)), where 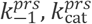 and *S*_lope_ are the pre-steady-state-like reverse first-order rate constants for the dissociation of ES into free E and S, a catalytic first-order rate constant, and a slope from the plot of the left-hand side of Eq. (22) versus [*S*_0_](1− exp. (-*k t*))/*k*, 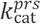 must be 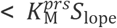 otherwise, the originating initial rates are only likely to be relevant where sQSSA is valid (*k*_cat_ > *K*_M_*k*_1_),

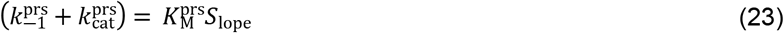

Note, however, that the slope is actually the second-order rate constant for the formation of ES. The result section gives better insight into the issues.

## 3. MATERIALS AND METHODS

### 3.1 Materials

#### 3.1.1 Chemicals

*Aspergillus oryzae* alpha-amylase (EC 3.2.1.1) and potato starch were purchased from Sigma-Aldrich, USA. Tris 3, 5—dinitrosalicylic acid, maltose, and sodium potassium tartrate tetrahydrate were purchased from Kem Light Laboratories in Mumbai, India. Hydrochloric acid, sodium hydroxide, and sodium chloride were purchased from BDH Chemical Ltd., Poole, England. Distilled water was purchased from the local market. The molar mass of the enzyme is ~ 52 k Da [21, 22].

### 3.2 Equipment

An electronic weighing machine was purchased from Wensar Weighing Scale Limited, and a 721/722 visible spectrophotometer was purchased from Spectrum Instruments, China; a *p*H metre was purchased from Hanna Instruments, Italy.

### 3.2 Methods

The enzyme was assayed according to the Bernfeld method [23] using gelatinised potato starch, whose concentration range was 0.3–3 g/L. Reducing sugar produced upon hydrolysis of the substrate at room temperature using maltose as a standard was determined at 540 nm with an extinction coefficient equal to 181 L/mol.cm. The duration of the assay was 3 minutes. A mass concentration of 2 mg/L of *Aspergillus oryzae* alpha-amylase was prepared in Tris-HCl buffer at *p*H *=*7.

#### 3.2.1 Determination of pseudo-first order rate constant, k and second order rate constant, k_1_

The determination of the pseudo-first order constant, *k*, for the utilisation of the substrate is as described elsewhere [24], with modification as follows: As in a manuscript under preparation, the result of the integration of a polynomial equation from the plot of initial rates *v*_i_ versus [*S*_0_] was fitted to the values of the former to give substrate concentrations that were < both the initial concentrations of the substrate and either *K*_d_ or *K*_M_. The new, but lower substrate concentrations were substituted into the polynomial equation to generate the corresponding lower velocities that were then used, as described in the literature [24], for the calculation of different values of *k*. The *k* values were then substituted into an equation as described in the literature [20] for the determination of the life span (*t*) of ES. Again, the value of *t* is substituted into Eq. (22) for the calculation of as described in the literature [20].

#### 3.2.2 Determination of ES dissociation constant, k_d_ and molar mass, M_3_ of potato starch

Equation (8) was applied in the determination of *k*_d_. The 2^nd^ equation for the determination of *M*_3_ is dependent on a reverse first-order rate constant for the dissociation of ES into free S and E. This is where Eq. (23) is relevant, and by being written as:

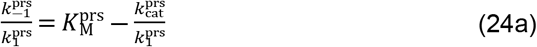

where 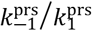 is = *k*_d_.

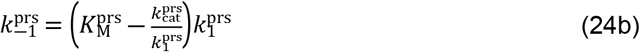

### 3.3 Statistical analysis

Assays were conducted in duplicate. The arithmetic mean of each initial rate was used to carry out double reciprocal plots and other plots. Micro-Soft Excel was explored for the determination of standard deviation (SD) where necessary.

## 4. RESULTS AND DISCUSSION

To begin with, it is imperative to note that whenever a plot of initial rates versus different substrate concentrations gives a negative coefficient of the leading term in a resulting polynomial, Michaelian kinetic characteristics are implicated; it may not be enough to guarantee the attainment of actual *K*_M_ and *V*_max_ at the asymptotic level if [*S*_0_] is not ≫ [*E*_0_]. The derived equation showed that if fitted to initial rates and plotted versus [*S*_0_] for an enzyme concentration of 2 mg/L, it gives a value that is < zero-order SC (Table 1). As Eq. (8) shows, the ES dissociation constant is directly proportional to the mass concentration, or molar concentration, if the molar mass of the enzyme is known, and to the molar mass of the substrate if it is known accurately. Thus, *K*_d_ is *ipso facto*, an established parameter. It is also amenable to experimental determination. The experimental values, obtained by a graphical approach (double reciprocal plot) and by calculation based on fitting a modified Michaelian equation (Eqs (18) and (19)) to initial rates at different [*S*_0_] and by Eq. (8), are, respectively, *K*_d_ ≈ 2.396 g/L and 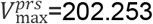 μM/min; *K*_d_ ≈ 2.407 g/L and 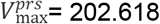 mM/min; and *K*_d_ = 2.48231 g/L.

**Table 1:** Experimental data-Independent and dependent variables.

|  |  |  |  |  |  |  |  |  |
| --- | --- | --- | --- | --- | --- | --- | --- | --- |
| $[S_0]/\text{g/L}$ | 0.3 | 0.6 | 0.9 | 1.2 | 1.5 | 1.8 | 2.0 | 3.0 |
| $v_i/\mu\text{M}/\text{min}$ | 22.514 | 40.5 | 55.2 | 67.4 | 77.8 | 86.7 | 91.9 | 113.3 |
| $K_M^{prs} \text{ (LWB)}/\text{g/L}$ | 2.396 | | | | | | | |
| $K_M^{prs} \text{ (Eqs (18 \& 19))}/\text{g/L}$ | $2.407 \pm 0.08 \text{ (} n=7 \text{)}$ | | | | | | | |
| $V_{\max}^{prs} \text{ (LWB) } / \mu\text{M}/\text{min}$ | 202.253 | | | | | | | |
| $k_{\text{cat}}^{prs} \text{ (LWB)}/\text{exp.}(+3)\text{min}$ | $\approx 5.259$ | | | | | | | |
| $V_{\max}^{prs} / \mu\text{M}/\text{min} \text{ (Eqs (18/19))}$ | $202.618 \pm 3.2 \text{ (} n=7 \text{)}$ | | | | | | | |
| $k_{\text{cat}}^{prs}/\text{exp.}(+3)\text{min Eqs.}(18\&19)$ | $\approx 5.268 \pm 0.083$ | | | | | | | |
| $K_d \text{ (Eq. (8))}/\text{g/L}$ | 2.482 | | | | | | | |
| $k_1/\text{exp.}(+4)\text{L/g. min}$ | 4.88 | | | | | | | |
| $k_{-1}(\text{LWB})/ \text{exp. (+5) min (Eq. (24b))}$ | $\approx 1.0975$ | | | | | | | |
| $k_{-1}/ \text{exp. (+5) min (Eqs (18\&19))}$ | $\approx 1.122$ | | | | | | | |
| $k_{-1}/ \text{exp. (+5) min (Eq.(8))}$ | $\approx 1.2112$ | | | | | | | |
| $M_3 \text{ (LWB)}/\text{exp. (6) g/mol.}$ | 62.296 | | | | | | | |
| $M_3 \text{ (Eqs (18 \& 19))}/\text{exp. (6) g/mol.}$ | 63.582 | | | | | | | |

$[S_0]$ , $K_d$ , and $V_{\max}^{prs}$ are the substrate concentration, ES dissociation constant, and pre-steady-state maximum velocity respectively; $k_{\text{cat}}^{prs}$ , $K_M^{prs}$ and $M_3$ are the pre-steady-state-like catalytic rate, pre-steady-state-like Michaelis-Menten constant, and molar mass of the substrate respectively. Based on Eq. (24a), $K_b$ is $\approx 2.288 \text{ g/L}$ using $K_M$ -like ( $K_M^{prs}$ ) result from Lineweaver-Burk (LWB) plot [27]; the value is $\approx 2.299 \text{ g/L}$ using $K_M^{prs}$ based on Eqs (18\&19).

The polynomial equation, generated from the plot of the initial rate versus [*S*_0_], is given as:

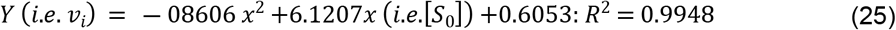

Equation (25) expresses a trend towards Michaelian kinetics due to the occurrence of a negative leading coefficient. This sQSSA relic contrasts with an almost perfect linear (*R*^2^ = 0.9993) relationship between calculated values of rate and calculated [*S*_0_] as described in Figure 1. This case is more characteristic of rQSSA. Further to this is the consideration of a situation in which the initial rate for [*S*_0_]_n_ is twice the initial rate for [*S*_0_]_n-1_; thus, with [*S*_0_]_1_ = 0.3 g/L and *v*_1_ = 2.251359 exp. (−5) M/min; [*S*_0_]_2_ = 0.6 g/L and *v*_2_ = 2 (2.251359) exp. (−5) M/min; [*S*_0_]_3_ = 0.9 g/L and *v*_3_ = 3 (2.251359) M/min exp. (−5) covering the first three data used for a plot, the unfolding result shows as expected, an equation of linear regression given as:

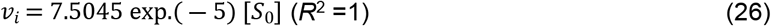

**Figure 1:**
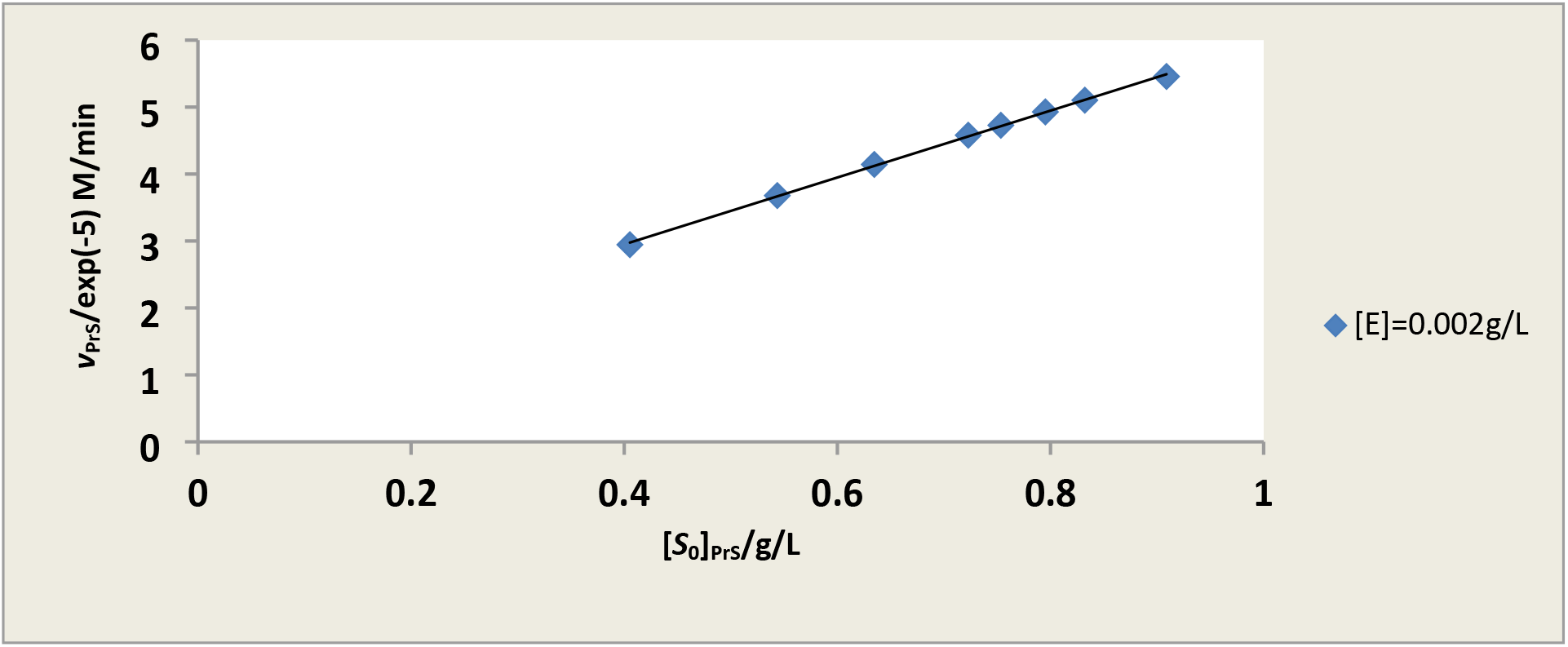
Experimental illustration of non-Michaelian linearised relation between calculated rates (*v*_prs_) from fitting polynomial equation to calculated [*S*_0_]_prs_ based on 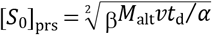 (manuscript under parallel preparation) where *M*_alt_, *v*, *t*_d_, α, and β are the molar mass of maltose, the product, velocity of hydrolysis, duration of assay, coefficient of the leading term, and coefficient of the term with unit power or exponent in the polynomial equation generated from the plot of initial rate versus [*S*_0_] (Eq. (24)).

However, it is important to note that only the first initial rate is directly experimental while the other two are calculated by multiplying the first rate by [*S*_n_]/[*S*_n-1_] for the purpose of illustration, a process not too different from simulation as applicable to well-known “high-reputation advanced publishers, FEBs, Elsevier, publisher of PNAS, Beilstein Journal, Biochemistry Journal (Oxford/Jn.), *etc.;* this research should not be an exception given that it is more of an experimental study with substantial theory. Dividing the slope by the molar concentration of the enzyme gives an SC-like value of 1951.17 L /g. min, which is < 2194.732 L/ g. min and ≈ 2188.645 L/g. min calculated from the table of values of and *K*_d_ (rewritten as) (Table 1); these values are, however, > the value (1298.414/g. min) obtained from the slope in Figure 1, a typical “rQSSA plot”.

As shown in this study, the *K*_d_ calculated on the basis of Eq. (24b) is different from the definite value obtained based on Eq. (8); this implies that it is not unlikely that different values of the experimental *K*_d_ can be obtained given different substrate concentration ranges for the same enzyme under the same assay condition. This is as long as the concentration range is < than the putative *K*_M_ value of the enzyme, and better still, it should be ≪ [*E*_0_] [6, 18]. A very important observation is that the 2^nd^ order rate constant *k*_1_ for the formation of the ES was determined and applied in the determination of the first-order rate constant for the dissociation of the ES into free E and S; different values (Table 1) are as a result of different values of 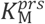 explored. The most important deductions are, however, the observation that the zero-order SC cannot be inferred from data points—the initial rates in particular—that either validate only rQSSA or partially validate sQSSA or by extension of the parameter domain that validates both rQSSA and sQSSA [25, 26]. The value of 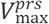 (≈124.198 mM/min) is based on the slope in Figure 1 and Eq. (10). This is a typical result that shows that the zero-order maximum velocity often inferred from Eq. (10) is inappropriate.

While noting a situation in which [*S*_0_]_n_*v*_n-1_ - [*S*_0_]_n-1_*v*_n_ is = 0, fitting a double reciprocal equation to such data gives a perfect straight line whose intercept is a small negative number while the slope is large; such a negative intercept does not show up if the three data points are part of the remaining five as shown below (Eq. (27d)). Table 1 shows the primary experimental data, the initial rates (average of duplicate studies, *n* = 2), and the corresponding concentrations of the substrate.

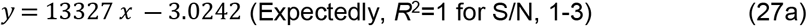

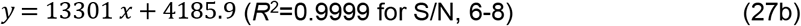

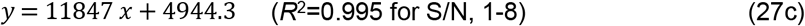

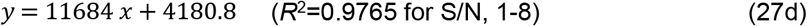

Equation (27d) is the outcome of the inclusion of the non-Michaelian initial rates (2 rates) that contributed to Eq. (26) in the double reciprocal plot. All plots, direct linear (or its reciprocal variant), and nonlinear regression, seem to mask the place and the role of the error introduced where the initial rates exhibit both rQSSA and a little bit of sQSSA validating attributes. In other words, where initial rates reproduce Eq. (26) and Figure 1 in any assay, nothing should be mentioned about sQSSA. Furthermore, Eqs (26) and (27a) are generally applicable where a single turnover event is desired, giving the impression that, in such a scenario, rQSSA is validly relevant because [*E*_0_] is ≫ the highest [*S*_0_] in the substrate concentration range chosen.

This study is very helpful considering the desire of biochemists, biophysicists, biochemical engineers, *etc*. to characterise the individual events of the catalytic cycle in the active site of enzymes. In this regard, individual events at the active site are easily isolated and studied without catalytic cycling, if single-turnover conditions are adopted [28]. In such a scenario, the substrate is saturated with enzyme ([*E*_0_] ≫ [*S*_0_]) so that all of the substrate will participate in the ‘single turnover’ [28]. This is the reason why the initial rate should always be directly proportional to [*S*_0_], as orchestrated by Eq. (10), and the demonstration of the implication of Eq. (10), represented by Eq. (26). The obvious is that in a true single turnover assay, the next higher initial rate will always be ([*S*_0_]_n_/[*S*_0_]_n-1_)-fold > than *v*_n-1_; this is the reason why in a real situation, apart from timing error and substrate depletion at the lower end of the substrate concentration range, there will be a small negative intercept (Eq. (27a)). This is clearly an expression of both Michaelian and sQSSA invalidity. This study used [*S*_0_] values, which are not very high compared to the [*E*_0_], though the latter is > all except 3 g/L of the insoluble gelatinised starch if the literature value of the molar mass of the insoluble potato starch is taken to be correct in the face of other values [29–31].

As depicted in Figure 2, there is a “far-right rQSSA” domain where it is impossible to infer any condition that validates the Michaelian equation and the associated sQSSA, as again illustrated by Eq. (26); this and Eqs (6 and 7) present the only means by which one can calculate the sub-zero-order maximum velocity, a peculiarity of a ‘single turnover’ catalytic activity whose conditions validate rQSSA. In this case, the molar masses of the substrate and enzyme, with known mass concentrations, must be known if the 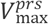 is to be calculated; otherwise, the slope indicated in Eq. (26) remains only a SC-like value. The value of 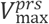 is ≈186.2724 μM/min. This value is clearly < than the values (Table 1) obtained from the LWB plot and Eqs (18 and 19). As shown in Figure 1 and Eq. (26), the values of 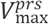 cannot be equal because of their different slopes; the result from Figure 1 is ≈ 123.964 μM/min. The point that cannot be ignored is that Eq. (1) cannot be used to calculate the *V*_max_ if the substrate concentration range is ≪ the known *K*_M_ of the enzyme whose concentration is either *≫* all concentrations of the substrate or ≈ [*S*_0_] [6, 18]. Any claim to the contrary, that *V*_max_ is known *a priori* for the determination of a mixed order (steady-state plus a near-zero-order state) *K*_M_, is invalid because all the [*S*_0_] values against which the pre-steady-state initial rates were plotted are < than the putative *K*_M_ value.

**Figure 2:**
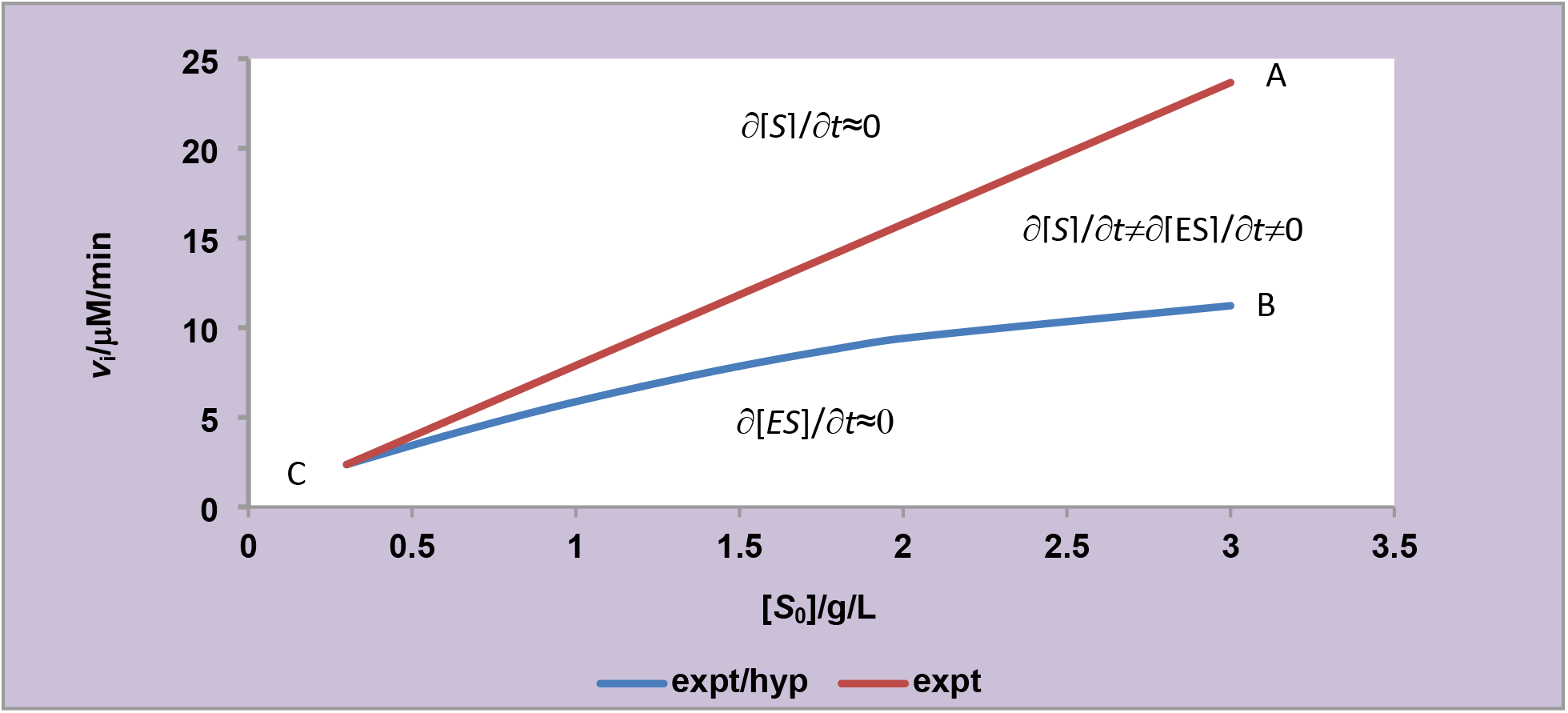
Graphical illustration of the domain of validity of different QSSA with the characteristic mathematical expression. Only the first data point in plot A, is experimental (expt.) while in plot B all the data points are experimental.

A clearer picture is obtainable considering Figure 2, where, anticlockwise from A, the condition that validates rQSSA with a higher concentration of the enzyme is the case [6, 18]. This is also a trend towards “single turnover” catalytic activity [28]. Clockwise, beginning from B, there is a higher tendency for the condition that validates both the Michaelian equation and the sQSSA. However, the view that “when both the sQSSA and rQSSA are invalid, the initial enzyme and substrate concentrations are comparable” seems to contradict the notion that [*S*_0_] needs not be ≫ [*E*_0_] for sQSSA to be valid and the claim in the literature [7] that the Michaelian equation and QSSA can still be valid if [*S*_0_]≈[*E*_0_]. Also contradicted is the notion of total QSSA (tQSSA), which is intended to extend the parameter domain for which both rQSSA and sQSSA could still be valid [25, 26]. In any case, what can be deduced from Figure 2 is that, as [*S*_0_] and [*E*_0_] tend towards equality, anticlockwise direction from B and clockwise direction from A make respectively the sQSSA and rQSSA less valid, but the transformation of the Michaelis-Menten equation can still be fitted to the initial rates as demonstrated with the experimentally generated equations, Eqs (27b-27d), unlike Eq. (27a).

The plot (Figure 2) shows mathematically that either only sQSSA and the Michaelian equation (line B and other lines that can be below it (the d[*ES*]/d*t*≈0 case)) or rQSSA (line A and any other lines above it (the d[*S*]/d*t*≈0 case)) if the first initial rate is half the next initial rate, and the corresponding concentrations of the substrate are such that the first is also half the next higher concentration of the substrate, which is peculiar to a single-turnover kinetics. Under the prevailing conditions between lines A and B, neither rQSSA nor sQSSA is fully validated; a shift of CA towards B through the middle invalidates it, while a shift of CB towards A invalidates it.

Figure 3 clearly demonstrates the strict relevance of rQSSA in this study, against the backdrop of the need to correctly specify the condition of the assay. This is apart from the physico-chemical aspects that influence the generated kinetic parameters. The linear regression of the initial rate, *v_i_*, versus the sub-*K*_M_ concentration of the substrates reflects one of such conditions that validate rQSSA and can only be used to determine the pre-steady-state (or rather, rQSSA) maximum velocity which is ≈ 135.348 mM/min based on Eq. (7) derived from Eq. (6), and the SC-like value is = 1417.65 L/g min.; such a value is not necessarily an exact value applicable to either a rQSSA (Eq. (6)) or sQSSA equation (linearised Michaelis-Menten equation). On the other hand, the inset showing LWBP does not necessarily produce the exact values of the kinetic parameters applicable to strict sQSSA, whose primary condition is the one that requires [*S*_0_] to be ≫ [*E*_0_] without which a true maximum velocity cannot be attained. The figure seems to confirm the claim in the literature that the Michaelian equation can still be valid if [*S*_0_] is ≈ [*E*_0_], or at least the former may not be ≫the latter.

**Figure 3:**
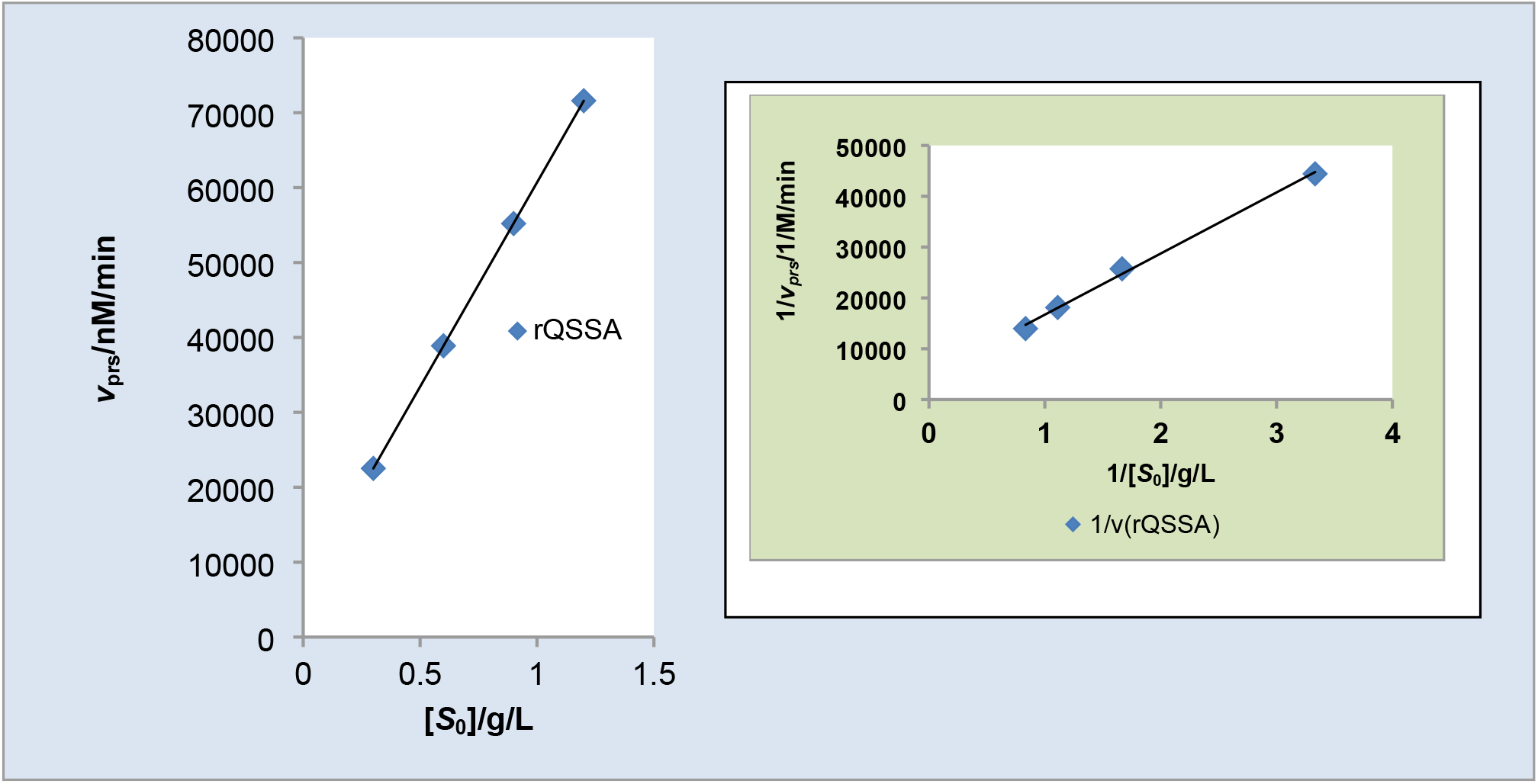
Plot illustrating how two set of similar initial rates can display both Michaelian kinetics (Lineweaver-Burk plot, LWBP, without negative intercept (*see the inset)* or polynomial with negative coefficient of the leading term) or sQSSA and rQSSA (a linear regression with *R*^2^ ≥ 0.999; in this case *R*^2^ is = 1). The *K*_M_-like value is ≈2.568 g/L; the *K*_d_ equivalent is = 2.5237 g/L and the molar mass of the starch based on Eq. (8) is ≈ 65.616 exp. (+6) g/mol.

The paper by Tzafriri and Edelman [26] exemplifies a scenario in which the rQSSA is applicable if the [*E*_0_] ≫ *K*_M_ and if the former is also as large as [*S*_0_]-the implication of such is depicted in Figure 3; but it must be made clear that *K*_M_ is [*E*_0_]-dependent. If the *K*_M_ remains at the substrate concentration at half maximum velocity, then any two or more different concentrations of the same enzyme under the same conditions must possess different *K*_M_ values, with the highest value referring to the highest concentration of the enzyme. Thus, different concentrations of the same enzyme under the same condition require different concentrations of the substrate for the attainment of maximum velocity (or for the orchestration of saturation phenomena) and consequently different values of *K*_M_. It is therefore obvious that in this study, where [*E*_0_] is > [*S*_0_], the condition relevant to rQSSA was very much the case; however, this is not to imply that there is no relic of sQSSA given the experimentally generated polynomial with a negative coefficient of the leading term. This notwithstanding, “reverse quasi-steady-state (rQSSA)” in which S is in a quasisteady state with respect to ES [1] characterises the main results obtained in this study, and it represents one of the few instances where quantitative effect as opposed to qualitative and pure mathematical analysis is carried out.

The preceding issues are further buttressed in Figure 4 which illustrates the domain where rQSSA and sQSSA are strictly valid (or upheld), and the domain in which QSSA as either rQSSA or sQSSA may be applicable or valid and beyond which neither may be valid. Line A (blue) which illustrates the domain of rQSSA validity, can become increasing valid if the concentration of the enzyme is increased for the same concentrations of the substrate [6, 17] leading to upward adjustment of line A; the converse is the case if the substrate concentration is decreased for the same concentration of the enzyme; line C (red) which illustrates the domain of sQSSA validity, can become increasing valid if the concentration of the enzyme is decreased for the same concentrations of the substrate leading to downward adjustment of line C (note that there could be upward adjustment if the concentrations of substrate is increased while the concentration of the enzyme is either decreased or remain the same and ≪ [*S*_0_]); the converse is the case if the substrate concentration is decreased for the same concentration of the enzyme; line B (green) has a dual representation of either conditions that validate rQSSA or sQSSA. Any increase in the concentration of the enzyme invalidates completely, the sQSSA, while any decrease in the concentration of the enzyme invalidates the rQSSA.

**Figure 4:**
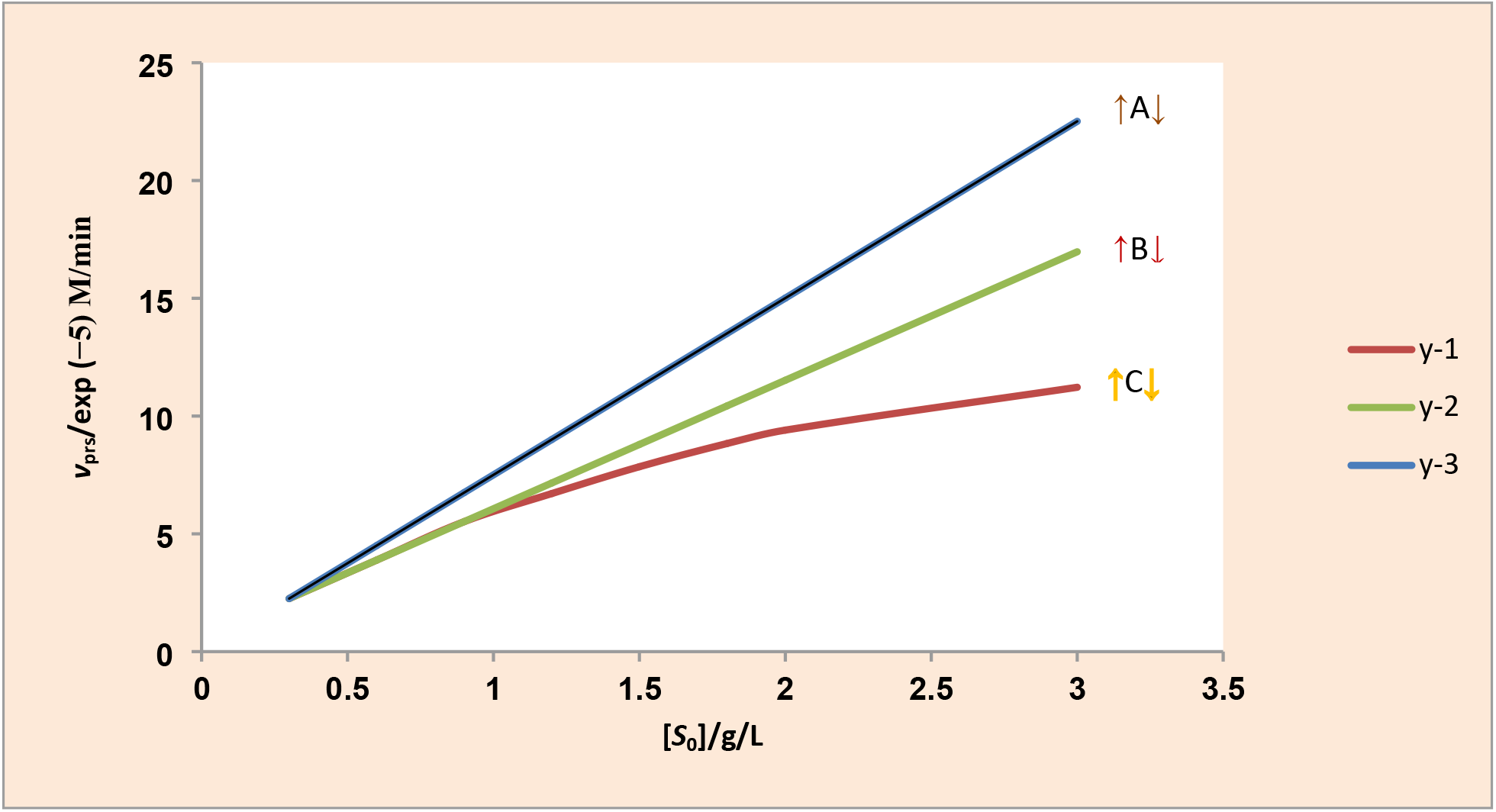
Experimental and simulational plots for illustrating the domain where strict rQSSA, strict sQSSA, and the domain in which QSSA as either rQSSA or sQSSA may be applicable or valid and beyond which neither may be valid.

Other kinetic parameters that are indirectly determined according to Eq. (22) are a reflection of the limit of the validity domain of sQSSA in favour of rQSSA, which has cognate kinetic constants such as the second order rate, *k*_1_, (for the formation of ES), and the reverse first-order rate, *k*_-1_, determined first based on Eq. (22) for the determination of *k*_1_, and second based on Eq. (23) for the determination of *k*_-1_ (see Table 1). This would have been impossible if the experimentally generated and simulated initial rates were applicable to sQSSA if the concentration of the enzyme was ≪ the concentration of the substrate. This has also made it possible to determine the molar mass of the insoluble potato starch if the label on the plastic container of starch purchased from Sigma is not faked by the distributor in the local major market. The molar mass is determinable given the following values (Table1): 62.296 exp. (+6) derived from LWB and 63.582 exp. (+6) g/mol. The calculated value based on Figure 3 (inset-LWB plot) is 65.616 exp. (+6) g/mol. The values compare with the cited literature values of 64.54 exp. (6) g/mol. [29]; however, a higher value of 77.3 exp. (+6) g/L [30] was also reported by the same author [31].

## 5. CONCLUSION

The equations which invalidate the assumption that *v*_i_ = *V*_max_ [*S*_0_]/*K*_M_ whenever [*S*_0_] is ≪ *K*_M_ were derived; the proposition that the equilibrium dissociation constant, *K*_d_, is strictly proportional to [*E*_0_] was confirmed with a derived equation; the calculated *K*_d_ value based on a rQSSA-derived equation could be > than the value obtained by graphical method; the molar mass of starch could be calculated from the derived equation; and it was shown graphically and mathematically that both sQSSA and rQSSA domains have a limit to their validity. The equation with which to calculate the second-order rate constant based on the condition that validates the rQSSA is not applicable to the sQSSA; a *K*_M_-like value that may be < the putative *K*_d_ value is considered for the assay of any concentration of an enzyme if the substrate concentration is ≪ actual *K*_M_. However, an assay with a substrate concentration range that is ≪ actual *K*_M_ may yield a *K*_M_-like value (≈2.569 g/L) that is > the putative *K*_d_ value of 2.482 g/L. The *K*_M_-like values in other situations are 2.396 and 2.407 g/L; the corresponding *K*_d_ values are, respectively, 2.288 and 2.299 g/L. The molar mass of insoluble potato starch ranges between 62.296 and 65.616 exp. (+6) g/mol. Contrary to the observation in the literature, in this study, mainly experimental and theoretical, the velocity, the initial rate, and the equations of the catalytic reaction have been employed on a number of occasions within the conditions for which they are valid and with reason why sQSSA may not be valid (reason: *k*_cat_ > *K*_M_*k*_1_ in such a scenario). A future study will explore the possibility of deriving a Lineweaver-Burk-like equation for the estimation of the molar mass of polymers like starch.

## ACKNOWLEDGEMENT

The management of the Royal Court Yard Hotel in Agbor, Delta State, Nigeria, is deeply appreciated for the supply of electricity during the preparation of the manuscript.

## REFERENCES

[1]. Segel LA, Slemrod M. Siam Rev. The quasi-steady-state assumption: A Case study in perturbation 1989; 31(3): 446–477. DOI: 10.1137/1031091.

[2]. Briggs GE, Haldane JB. A note on the kinetics of enzyme action. Biochem. J. 1925; 19: 338–339. DOI: 10.1042/bj0190338.

[3]. Savageau MA. Biochemical systems analysis: I. Some mathematical properties of the rate law for the component enzymatic reactions. J. Theor. Biol. 1969; 25: 365–369. DOI: 1016/s0022-5193(69)80026-3

[4]. Bakalis, E, Kosmas M, Papamichael EM. Perturbation theory in the catalytic rate constant of the Henri–Michaelis–Menten enzymatic reaction. Bull. Math. Biol. 2012; 1–15. DOI 10.1007/s11538-012-9761-x.

[5]. Van Slyke DD, Cullen GE. The mode of action of urease and of enzymes in general. J. Biol. Chem. 1914; 19: 141–180. DOI: 10.1016/s0021-9258(18)88300-4.

[6]. Schnell S, Maini PK. Enzyme Kinetics at High Enzyme Concentration. Bull. Math. Biol. 2000; 62: 483–499. DOI:10.1006/bulm.1999.0163.

[7]. Schnell S. Validity of the Michaelis-Menten equation – steady-state or reactant stationary assumption: that is the question. FEBS J. 2014; 281: 464–472. DOI:10.1111/febs.12564.

[8]. Schnell S, Maini PK. A Century of Enzyme Kinetics: Reliability of the *K*_M_ and *V*_max_ Estimates. Comments on Theoretical Biology, 2003; 8: 169–187. DOI: 10.1080/08948550390206768.

[9]. Cornish-Bowden A, Endreny L. Fitting of enzyme kinetic data without prior knowledge of weights Biochem J. 1981; 193: 1005–1008. DOI: 10.1042/bj193105.

[10]. Ritchie RJ, Pyran T. Current statistical methods for estimating the *K*_M_ and *V*_max_ of Michaelis-Menten kinetics. Biochem. Edu. 1996; 24 (4), 196–206. DOI: 10.1016/50307-4412(96)00089-1.

[11]. Marasović M, Marasović T, Miloš M Robust nonlinear regression in enzyme kinetic parameters estimation. J. Chem. 2017; 1–13. DOI: 10.1155/2017/6560983.

[12]. Matyska L & kovář J. Biochem. J. Comparison of several non-linear-regression methods for fitting the Michaelis-Menten equation 1985; 231: 171–177. DOI: 10.1042/bj2310171.

[13]. Nelatury SR., Nelatury CF, Vagula MC. (2014) Parameter estimation in different enzyme reactions. Adv. Enz. Res. 2014; 2: 14–16. DOI: 10.4236/aer.2014.21002.

[14]. Johnson K.A. New standards for collecting and fitting steady-state kinetic data Beilstein J. Org. Chem. 2019; 15: 16–29. DOI: 10.3762/bjoc.15.2.

[15]. Udema II. Rate constants are determinable outside the original Michaelis–Menten mathematical formalism wherein the substrate concentration range is approximately 1.6 to 4.8 times enzyme concentration: A pre-steady-state scenario and beyond. World J. Adv. Res. Rev. 2022; 16(01): 350–367. DOI: 10.30574/wjarr.2022.16.1.0989.

[16]. Udema II. Derivation of steady-state first-order rate constant equations for enzyme-substrate complex dissociation, as well as zero-order rate constant equations in relation to background assumptions. GSC Biol. Pharm. Sci. 2022; 03: 175–189 DOI:10.30574/gscbps.2022.21.3.0482.

[17]. Schnell S, Maini PK. Enzyme kinetics far from the standard quasi-steady-state and equilibrium approximations. Math. Comput. Model. 2002; 35: 137–144. DOI: 10.1016/S0895-7177(0100156-x).

[18]. Borghans, JAM, de Boer RJ, Segel LA. Extending the quasi-steady state approximation by changing variables. Bull. Math. Biol. 1996; 58: 43–63. DOI:10.1007/BF02458281.

[19]. Udema II. Alternative equations and “pseudo-statistical” approaches that enhance the precision of initial rates for the determination of kinetic parameters. BioRxiv preprint. 2023; 1–23. DOI: 10.1101/2023.01.16.524223

[20] Udema II. Derivable equations and issues often ignored in the original Michaelis-Menten mathematical formalism Asian J. Phys. Chem. Sci. 2019; 4: 1–13. DOI: 10.9734/ajopacs/2019/7i430101.

[21]. Udema II. Derivation of kinetic parameter dependent model for the quantification of the concentration and molar mass of an enzyme in aqueous solution: A Case study on *Aspergillus oryzae* α-amylase. Journal of Scientific Research & Reports 10(3): 1–10. DOI: 10.9734/JSRR/2016/24321.

[22]. Sugahara M, Takehira M, Yutani K. Effect of heavy atoms on the thermal stability of alpha-amylase from Aspergillus oryzae. Plos One. 2013; 2: 1–7. Doi:10.137/journal.pone.0057432

[23]. Bernfeld P. Amylases, alpha and beta. Methods. Enzymol. 1955; 1: 149–152. DOI: 10.1016/0076-6879(55)01021-5.

[24]. Udema II, Onigbinde AO. The experimentally determined velocity of catalysis could be higher in the absence of sequestration. Asian. J. Res. Biochem. 2019; 5(4): 1–12. DOI: 10.9734/AJRB/2019/v5i430098.

[25]. Tzafriri AR. Michaelis–Menten kinetics at high enzyme concentrations. Bull. Math. Biol. 65: 2003; 1111–1129. DOI: 1016/S0092-8240(03)00059-4.

[26]. Tzafriri AR, Edelman ER. Quasi-steady-state kinetics at enzyme and substrate concentrations in excess of the Michaelis–Menten constant. J. Theor. Biol. 2007; 245: 737–748. DOI:10.1016/j.jtbi.200612.005.

[27]. Lineweaver H, Burk D. The determination of enzyme dissociation constants. J. Am. Chem. Soc. 1934; 3: 658–666. DOI:10/1021/ja01318a036.

[28]. Sassa A, Beard W.A., Shock D.D., Wilson S.H. steady-state, pre-steady-state, and single-turnover kinetic measurement for DNA glycosylase Activity. J. Vis. Exp. 2013; (78): 1–9. DOI: 10.3791/50695.

[29]. Lii CY, Lia CO, Stabińsk L, Tomasik P. Effect of corona discharge on granular starch J. Food Agric. Environ. 2003a; 1: 143–149.

[30]. Lii CY, Lia CO, Stabińsk L, Tomasik P. Effect of hydrogen, oxygen, and ammonia low-pressure glow plasma on granular starches. Carbohydr. Polym. 2002b; 49: 449–456. DOI: 10.1016/S0144-8617(01)00351-4.

[31]. Tomasik P. Specific chemical and physical properties of potato starch. Food. 2009; 9: 45–46.

